# Beliefs about a widely popularised human-wildlife conflict are largely unreflective of local experiences and ecological data in wild red foxes, *Vulpes vulpes*

**DOI:** 10.64898/2025.12.06.692778

**Authors:** Kristy A. Adaway, Charlotte R. Hopkins, Carl D. Soulsbury, F. Blake Morton

## Abstract

Human-driven environmental changes, such as urbanisation, are increasing human-wildlife interactions, creating potential for conflict. Public beliefs and experiences with local wildlife are central to the success of place-based conservation initiatives, such as urban rewilding, because they impact public attitudes and tolerance of those species within neighbourhoods. Few studies, however, have examined whether people’s beliefs align with their real-life observations of wildlife within their local area and, more importantly, whether (or how) their beliefs might impact public attitudes independently of local experiences with those animals. The current study addressed this gap by assessing whether public beliefs about a conspicuous human-wildlife conflict – bin-raiding behaviour – aligned with residents’ own local experiences with a familiar and widespread urban carnivore, the red fox (*Vulpes vulpes*). The study also examined whether beliefs about bin-raiding predicted attitudes towards foxes even when people lacked firsthand observations of such behaviour within their area. Beliefs and attitudes towards foxes were evaluated using a nationwide questionnaire administered to 1,275 participants in the United Kingdom. Ecological surveys were also conducted in the city of Hull, UK, to compare public attitudes and beliefs (N= 248 households) with objective data on outdoor bin disturbances (N=4,239 bins) from the same area. Nationally, most respondents (59.3%) believed that foxes raid bins despite significantly fewer of these people (21.7%) having witnessed it firsthand within their neighbourhood. In Hull, <1% of bins showed evidence of disturbance even though significantly more residents (16.9%) believed foxes raided bins than the number of residents who had actually seen it happen. Among national and local Hull residents with no local experience observing bin-raiding foxes, these beliefs were nevertheless associated with more negative attitudes towards them. Together, these findings highlight the importance of taking into account other factors beyond local experiences (e.g., socio-cultural norms) when interpreting local reports of human-wildlife conflict, especially for place-based conservation.

## Introduction

Human-driven environmental changes, including urbanisation and climate change, are transforming ecosystems and increasing interactions between wildlife and people (Schell et al., 2021; Abrahms et al., 2023). These interactions can provide opportunities for coexistence, but they can also lead to human-wildlife conflict, defined in terms of negative interactions that adversely affect one or both parties (Conover, 2002; Abrahms, 2021; Conover & Conover, 2022). For humans, these impacts can include vehicle collisions, economic losses (e.g., crop, livestock, or property damage), and risks to health (e.g., zoonoses, injury, or death), along with psychological stress and reduced well-being (Barua et al., 2013; Soulsbury & White, 2015; Nyhus, 2016; Schell et al., 2021; Conover & Conover, 2022; Bombieri et al., 2023). Collectively, such interactions amount to costs of billions of US dollars each year (Nyhus, 2016; Conover & Conover, 2022). Human-wildlife conflict can also impact public attitudes and reduce tolerance towards individual species (Gillingham & Lee, 2003; Hazzah et al., 2009; Liu et al., 2011; Kifle, 2021; Hobson et al., 2024; Puri et al., 2024), which may undermine conservation efforts aimed at supporting wildlife (Hazzah et al., 2009; Kifle, 2021; Puri et al., 2024). Such challenges are particularly important for place-based conservation strategies where success depends not only on ecological factors, but also on local public support (Sifuna, 2010; Vasudeva et al., 2021; Malviya et al., 2022; Auster et al., 2025). Understanding how attitudes form, and how they are shaped by direct experiences versus other factors (e.g., socio-cultural norms), is therefore critical for fostering coexistence and ensuring the long-term success of conservation initiatives in human-dominated landscapes.

One example of how public attitudes can impact place-based conservation is rewilding, which aims to restore specific ecosystems that have been significantly altered by human activities by re-establishing natural processes, thereby creating self-sustaining and resilient environments (Carver et al., 2021). In urban contexts, rewilding efforts typically focus on enhancing biodiversity by increasing native vegetation, restoring natural habitats, and improving habitat connectivity, with a goal of creating conditions that allow wildlife to recolonise and thrive in the future (Pettorelli et al., 2022). However, human-wildlife conflict already occurs in many urban settings (Soulsbury & White, 2015; Nyhus, 2016; Basak et al., 2020; Schell et al., 2021; Basak et al., 2023; Moesch et al., 2024), and rewilding could amplify these interactions by increasing the presence and visibility of wild animals (Pettorelli et al., 2022; Rosenkrantz, 2023). Closer, more frequent interactions raise the potential for conflict, which may lead to more negative attitudes and efforts to control or remove species, ultimately undermining the goals of rewilding (Pettorelli et al., 2022). Public support is therefore essential, but crucially, opposition may stem not only from actual conflict with wildlife, but also by *perceived* conflict within the local area, that is, people’s beliefs that certain species are problematic even when no concrete harm has been experienced.

Previous research suggests that people’s beliefs about wildlife conflict may not always reflect the level of threat or damage actually displayed by those animals within their local area (Gillingham & Lee, 2003; Richard et al., 2005; Hazzah et al., 2009; Dickman, 2010; Echegaray & Vilà, 2010; Dickman et al., 2014; Ballejo et al., 2020; Bridge & Harris, 2020; Wilkinson et al., 2021; Songhurst, 2023; Iranzo et al., 2025; Vela-Vargas et al., 2025). Beliefs – whether accurate or not – can feed into broader attitudes towards a species (Tarrant et al., 2016; Baker et al., 2020; Kimmig et al., 2020). When beliefs exaggerate conflict, they tend to foster more negative attitudes, which can directly undermine human-wildlife coexistence by encouraging persecution (Richard et al., 2005; Hazzah et al., 2009; Echegaray & Vilà, 2010; Ballejo et al., 2020). Echegaray and Vilà (2010) found that free-ranging dogs could potentially be responsible for some livestock attacks in the Basque Country, Spain, despite grey wolves (*Canis lupus*) being widely blamed. Similarly, Ballejo et al. (2020) found that farmers in Patagonia believed scavenging birds frequently attacked livestock, yet field data showed little evidence of such predation. In both cases, public misconceptions shaped negative attitudes and ultimately lethal control of wolves and birds, illustrating how perceived local conflict can drive management actions that affect biodiversity (Echegaray & Vilà, 2010; Ballejo et al., 2020). However, despite evidence that perceived local conflict can contribute to wildlife persecution, wildlife management typically relies heavily on subjective reports of conflict by residents, largely due to limited resources (Dickman, 2010). By comparison, there are relatively few examples where these reports have been rigorously compared to direct observations of animal behaviour from the *same* location, making it difficult to distinguish localised experiences with conflict versus other factors, such as broader generalisations about a species. This gap highlights the urgent need for more research to distinguish real versus perceived human-wildlife conflict when interpreting local reports of human-wildlife conflict. Such distinctions are particularly crucial in light of global urbanisation and the growing popularity of place-based conservation strategies, such as urban rewilding, which increase the likelihood of encounters between wildlife and people.

To address this gap, the current study investigated the extent to which public beliefs about a wild urban carnivore, the red fox (*Vulpes vulpes*), are underpinned by personal local observations of them, and how beliefs relate to broader public attitudes towards the species even when people lack firsthand experience observing their behaviour within their neighbourhood. Foxes are a highly adaptable species inhabiting many habitat types throughout Europe, Africa, Asia, North America, and Australia (Harris, 1981; Contesse et al., 2004; Soulsbury et al., 2010; Plumer et al., 2014; Scott et al., 2014; Gil-Fernández et al., 2020; Morton et al., 2023). As the most widespread terrestrial carnivore on the planet (Macdonald & Reynolds, 2004; Soulsbury et al., 2010), human encounters with foxes are becoming more prevalent in light of on-going urbanisation (Plumer et al., 2014; Scott et al., 2014; Basak et al., 2022). Although foxes already thrive in urban landscapes, recent rewilding efforts aiming to enhance urban habitat quality and connectivity could result in further close human-fox encounters by helping foxes thrive and expand into other neighbourhoods (Baker & Harris, 2007; Bateman & Fleming, 2012). Thus, to promote coexistence and ensure that urban rewilding initiatives are successful, there is a pressing need to better understand the drivers of public attitudes and beliefs towards the species.

Foxes are often portrayed in popular culture as prolific bin-raiders, i.e., animals that rummage through rubbish bins in search of food (e.g., The Newsroom, 2013; Dubuis, 2014; Reynolds, 2025). Bin-raiding behaviour, and the mess it is believed to cause, is frequently cited as a key reason for public disapproval of urban foxes (Harris, 1981; Baker et al., 2020; Brand & Baldwin, 2020). However, recent ecological research suggests that this reputation may be overstated (Harris, 1981; Contesse et al., 2004; Plumer et al., 2014; Morton et al., 2023). For example, Morton et al. (2023) tested foxes’ interactions with food-related objects to simulate the kinds of barriers presented by rubbish bins, and found that only 12.5% of foxes extracted food from the objects after two weeks. Previous research suggests that bin-raiding may be restricted to specific individuals or populations rather than being typical of foxes more generally (Harris, 1981; Contesse et al., 2004; Morton et al., 2023). Further, it is possible that beliefs about foxes raiding bins stem from outdated experiences. For example, in the 1980s, UK councils began replacing traditional dustbins with more secure wheelie bins (Chappells & Shove, 1999), which are harder for foxes to access, and some sources report that the already low levels of bin-raiding by foxes further declined after this change (Baker et al., 2004; Baker et al., 2006). Taken together, beliefs about bin-raiding may be shaped by factors beyond people’s more recent experiences with foxes within their immediate area. However, without comparing public beliefs and experiences to actual bin disturbances from the same locations, such dynamics are impossible to disentangle.

Given that previous ecological studies suggest foxes may not raid bins as often as people think (Harris, 1981; Contesse et al., 2004; Plumer et al., 2014; Morton et al., 2023), the current study hypothesised that respondents would be more likely to say they believe foxes raid bins compared to the number of people who report having directly observed such behaviour within their neighbourhood. The study also hypothesised that beliefs about bin-raiding would predict attitudes towards foxes among people who lacked firsthand local experiences.

## Methods and Materials

### Ethical approval

The study was approved by the Faculty of Health Sciences Ethics Committee of the University of Hull (FHS 23-24.30 & FHS 24-25.008) and followed the research guidelines of the British Psychological Society.

### Survey method and design

#### Measuring fox-related attitudes, experiences, and beliefs

Data on public fox-related attitudes, experiences, and beliefs were collected using an online questionnaire administered at two spatial levels – at the national level and at a more local level within the city of Hull, UK. The national-level analysis was conducted across urban and rural areas throughout Scotland, England, and Wales between March and April 2023. The local-level analysis was conducted in Hull between January and February 2025.

For the national survey, participants were recruited via the commercial survey provider Survation. For the local survey, the survey was advertised through the use of fliers delivered to 8,074 households within the HU88 and HU54 postal areas via the Royal Mail leaflet drop service (Royal Mail, 2025). These areas cover large residential areas and aligned with areas where bin surveys were conducted.

The national survey was initially designed to test whether providing information about fox psychology influenced people’s attitudes towards foxes, and whether this effect varied depending on the delivery method (e.g., press release or video) (Morton et al., 2024). The survey consisted of three parts: Part A captured demographic information and participants’ experiences with fox bin-raiding. Part B presented information about fox psychology or ecology. Part C assessed attitudes and beliefs about foxes using a 24-item Likert scale (Morton et al., 2024). Participants attitudes were not significantly influenced by either the information or the format used in Part B (Morton et al., 2024); thus, their responses from Parts A and C were used for the current study. Full details of the survey can be found in Morton et al. (2024).

For both questionnaires, participant’s beliefs about bin-raiding were evaluated by asking on 7-point Likert scale to what extent they agreed to the following statement: “Foxes try to get food from peoples’ outdoor rubbish bins”. People who scored 1-3 on the scale were classified as “disagreeing” that foxes raided bins, while scores of 4 were classified as “neutral”, and scores of 5-7 were classified as “agreeing”. People’s personal observations of fox bin-raiding from their local area were evaluated through two questions (Questions 13 and 16 in Part A of the national survey, and 12 and 19 in Part A of the local survey) asking whether, at their current place of residence, they had ever witnessed (in person or via camera) a fox eating rubbish from their own outdoor bin or a public bin in their neighbourhood. Responses to these two questions were combined into a single binary measure of local experience with foxes eating from a bin and were classified as yes (1) if they had personally seen a fox eating from a bin, and no (0) if they had not seen a fox eating from a bin. For the survey administered in Hull, participants were specifically asked whether they had witnessed bin-raiding foxes within the last year (whereas the national survey did not), as the intention was to align these responses with the ecological surveys that were conducted in the same areas during the previous year (see below).

Previous research has shown that inventories to evaluate general attitudes towards foxes are comprised of just a single factor, with people’s scores ranging from relatively positive to negative, respectively (Morton et al., 2024). Thus, in the current study, a single composite score was calculated by averaging responses to positive items, subtracting the average of negative items, and producing a score from -6 (very negative) to +6 (very positive). For the national analysis, this was based on 22 items (Questions 2 to 9 and 12 to 25) from Part C of the original Morton et al. (2024) questionnaire. For the local analysis, general attitudes were based on 18 Likert items (Questions 1 to 5 and 8 to 20) from Part B of the survey (Supplementary Materials).

#### Ecological surveys on outdoor bin disturbances

Ecological surveys were conducted concurrently with the questionnaire administered to people within Hull, allowing us to directly compare people’s attitudes and beliefs about foxes to objective data on outdoor bin disturbances within the same neighbourhoods. Specific streets included in the ecological survey were determined using a crosshair line sampling method, allowing for a representation of different residential (terrace housing estates, semi-detached housing estates), commercial and industrial areas as seen in other studies (Bjerke & Østdahl, 2004; Contesse et al., 2004). A list of the post codes that fell along the crosshair lines was compiled, and the Hull City Council website was used to determine when the local council attended each postcode to empty their outdoor waste bins (Hull City Council, 2023).

Household waste bins were surveyed as they were left out on the public street to be emptied by the local council on their waste collection days between the 8th of January and the 16th of February 2024. Bin surveys began in the early morning, around 7am just before first daylight and continued until the council arrived within the sampling area, around 11am. On these days, households are required to leave their bins on the street for collection by 7am, and many residents place them outside on the night of the previous day. Bin surveys were conducted on foot by walking the streets along the crosshair lines and taking high-quality photos and videos of the bins. Upon completion of the street surveys, ExifTool (Harvey, 2024) was used to extract meta-data, including the date, time, and GPS coordinates for each individual photograph or video taken of bins along the crosshair lines.

The photographic and video data were systematically coded using predefined criteria to visually determine whether the contents of the bin were easily accessible to animals, and whether there was any evidence of the bin being disturbed by something (**Table 1.1**). Foxes are not the only urban species with the ability to extract food from bins, e.g., dogs, cats and badgers (Harris, 1981), and it was not possible to identify fox-specific bin disturbances (Contesse et al., 2004). Therefore, to account for ambiguous cases (e.g., bin bags that could have been split open due to their contents), two additional measures of “disturbed bins” were coded: a conservative measure, including only bins highly likely to have been disturbed by an animal (possibly a fox), and a less conservative measure, which included bins with more ambiguous signs (**Table 1.1**). Inter-observer reliability (IOR) tests were then conducted to assess the degree of agreement between the original coding and that of independent coders who reviewed the data.

**Table 1.1.**
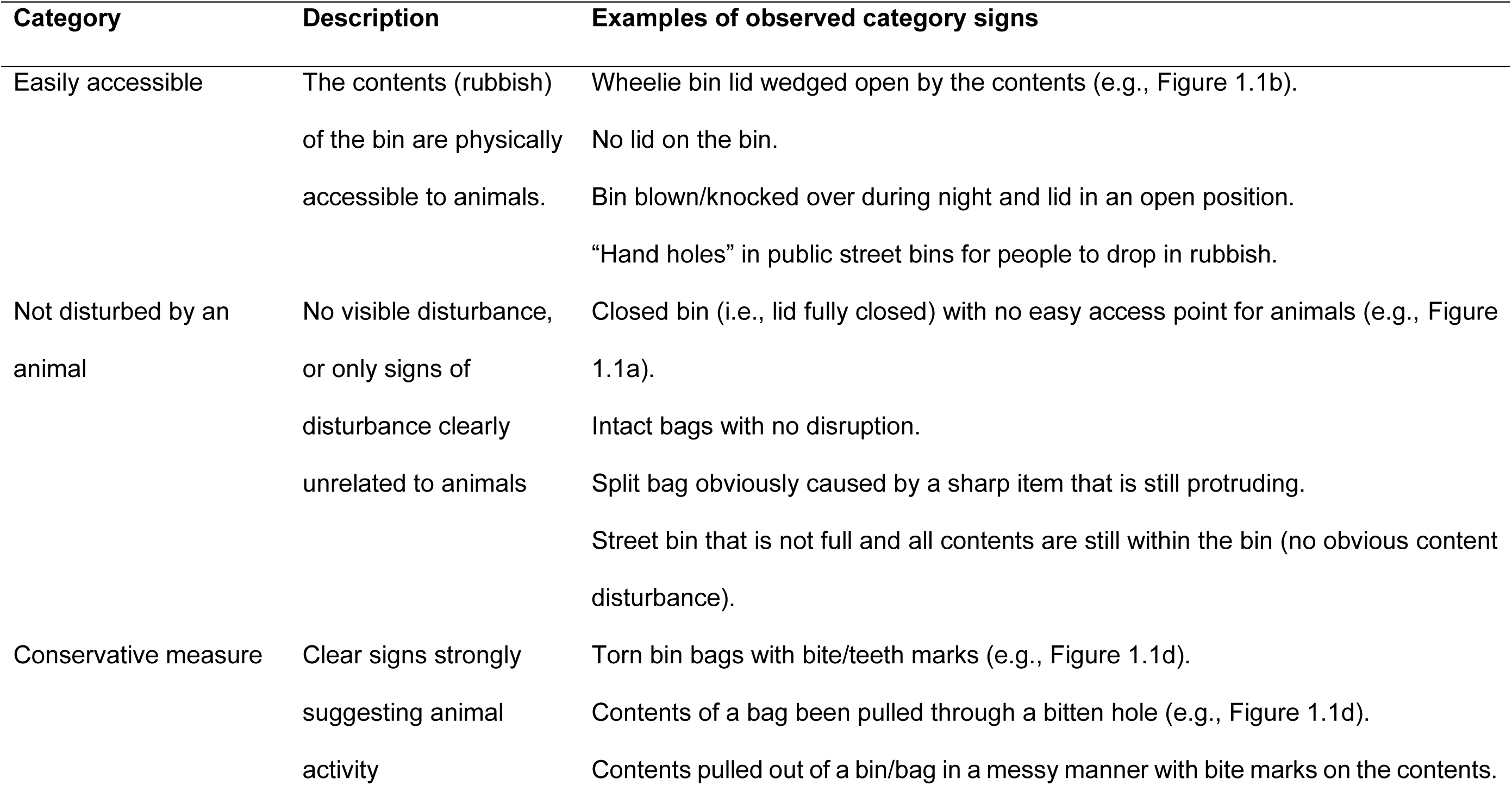
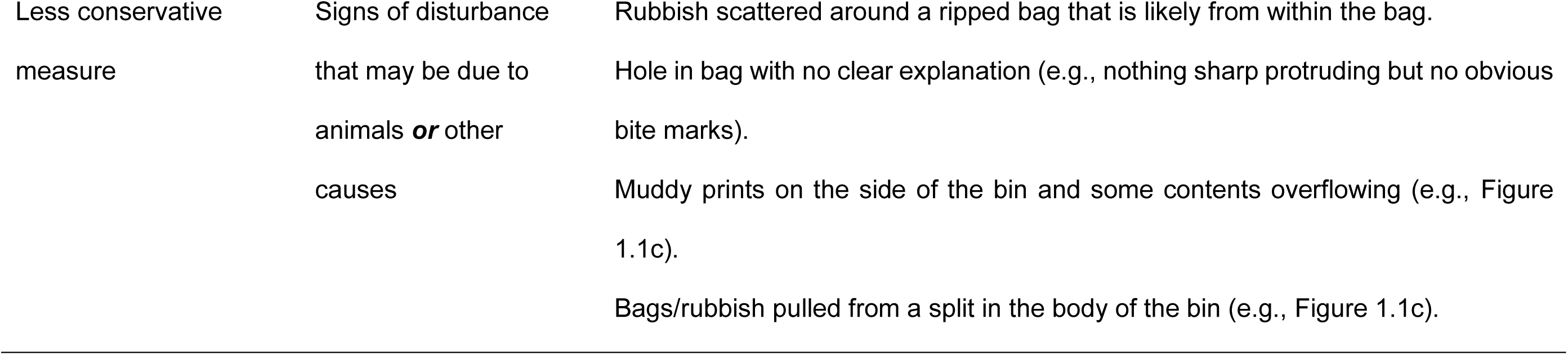
Coding criteria for bins from ecological surveys in Hull, UK.

Within the city of Hull, three main types of outdoor household bins are used by local residents for food waste, including 1) grey/black wheelie bins for food and general household waste, 2) brown wheelie bins for food and garden waste, and 3) green caddies for food waste only. Bin types vary by area, and many households have two types collected on alternating weeks. Thus, to ensure all bin types were surveyed, areas with multiple bin types were revisited on the alternate collection week, with only the previously un-surveyed bins recorded. Public spaces that fell along the crosshair lines (e.g., parks, cemeteries), and commercial and industrial areas were also surveyed. In these areas, bins included public street bins owned by the local council, which typically contain mixed litter such as food packaging and drinks containers; large commercial dumpsters, which generally contain bulk refuse, including food waste, from shops, restaurants, and workplaces; smaller, differently coloured wheelie bins provided for public rubbish disposal in town centres; and occasional bin bags placed directly on the ground, usually holding household waste. However, there was no publicly available information regarding when these bins were emptied (i.e., when they were likely to be at their fullest). As a result, these areas were surveyed at the same time as the nearest residential area.

**Figure 1.1.**
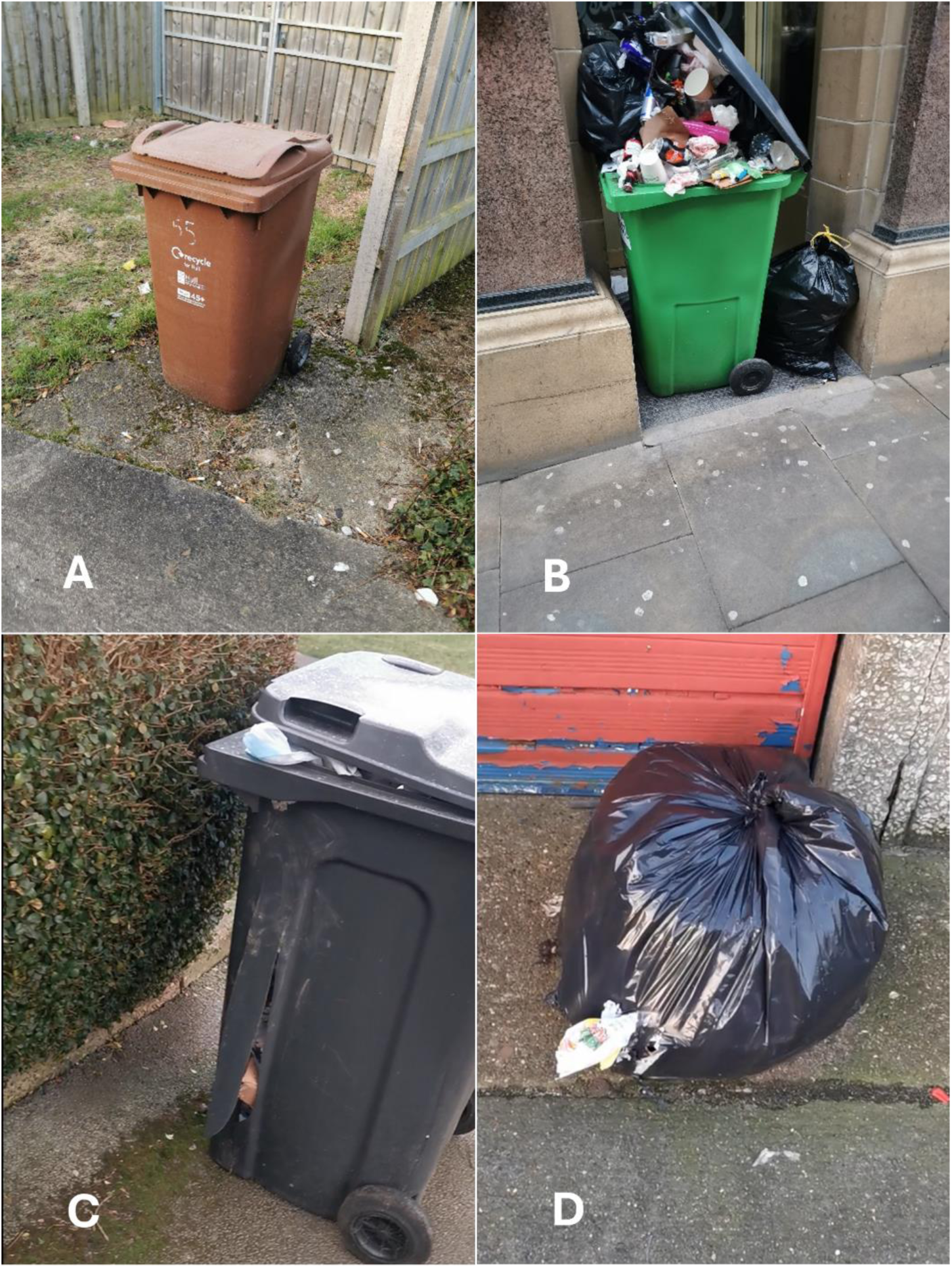
Photos of bins taken during the ecological surveys in Hull, UK, including a) a closed brown wheelie bin, b) a public wheelie bin in town centre overflowing with waste but without evidence of animal disturbance, c) a black wheelie bin with possible signs of animal disturbance (i.e., muddy prints and contents potentially pulled through split in bin), and d) a black bin bag with signs of animal disturbance (i.e., bite marks in bag and contents pulled through the hole).

### Statistical analyses

All statistical analyses were conducted using R version 4.3.3 (R Core Team, 2024). Two-Proportion Z-tests were used to determine whether there was a significant difference in the proportion of respondents who had witnessed a fox eating from bin (either their own bin, or a public bin owned by their local council) and the proportion of respondents that agreed that foxes raid bins.

To isolate the effect of belief about fox bin-raiding on attitudes independent of local personal experience, we focused analyses on participants who had not witnessed local bin-raiding. Among these respondents, Kruskal-Wallis rank sum tests were used to compare participants’ attitudes towards foxes based on whether they agreed, disagreed, or were neutral about the belief that foxes raid bins. To ensure that the outcome measure reflected general attitudes rather than overlapping with the predictor, two Likert items specifically addressing beliefs about foxes raiding bins (including the item used to categorise participants’ beliefs) were excluded from the composite attitude scores. Excluding respondents with local bin-raiding experience allowed us to examine whether holding the belief that foxes raid bins was associated with more negative attitudes even when participants had no direct local experience with bin-raiding. Post-hoc Dunn’s tests with Benjamini-Hochberg correction for multiple comparisons were conducted using the FSA package to identify significant differences (Ogle et al., 2023).

Cohen’s kappa tests were conducted to determine the inter-observer reliability agreement for the ecological surveys conducted in Hull. When considering which bins were easily accessible to animals, there was excellent inter-observer agreement (k > 0.75) between K. A., and an independent coder (C. B.) who coded 100% of the data (Table S4.1). When considering which bins were highly likely to have been disturbed by an animal that could have been a fox (i.e., the conservative measure), there was excellent agreement (k > 0.75) between K. A. and a second independent coder (K. S.) who coded 100% of the data (Table S4.2). When considering which bins were “potentially disturbed by an animal” (i.e., the non-conservative measure), there was good inter-observer agreement (k = 0.63) between K. A., and a third independent coder (D. J.) who coded 50 samples of the data (Table S4.3). All Cohen’s kappa test were conducting using the thresholds developed by Cicchetti and Sparrow (1981) and using the IRR package (Gamer & Lemon, 2019).

Maps were created using Base map data from OpenStreetMap contributors (2025) under the Open Database Licence (ODbL). Boundary outlines for maps were obtained from Planning Data (2024) under the Open Government Licence v3.0. Figures were produced in R using the package ggplot2 (Wickham, 2016).

## Results

### Public attitudes, beliefs, and experiences with foxes

In total, 1,373 people responded to the national public survey. However, 98 respondents were excluded, including nine people because they were unable to correctly identify a fox, and a further 89 people because their prior bin-raiding experiences could not be determined due to their failure to answer the question. The remaining 1,275 respondents ranged in age from 18 to 93 years (median = 58, interquartile range = 38 – 69). Of these people, 756 (59.3%) participants agreed with the statement that foxes try to get food from peoples’ outdoor rubbish bins, while 140 (11%) and 379 (29.7%) participants disagreed or were neutral, respectively. In terms of their personal observations of fox bin-raiding within their local area, 277 (21.7%) people said that they had witnessed bin-raiding, while the remaining 998 (78.3%) people said that they had not. The proportion of respondents who agreed that foxes raid bins was significantly higher than the proportion who reported having personally witnessed fox bin-raiding within their local area (Two proportion Z test: X^2^_1_= 371.8, p < 0.001; **Figure 1.2**).

**Figure 1.2.**
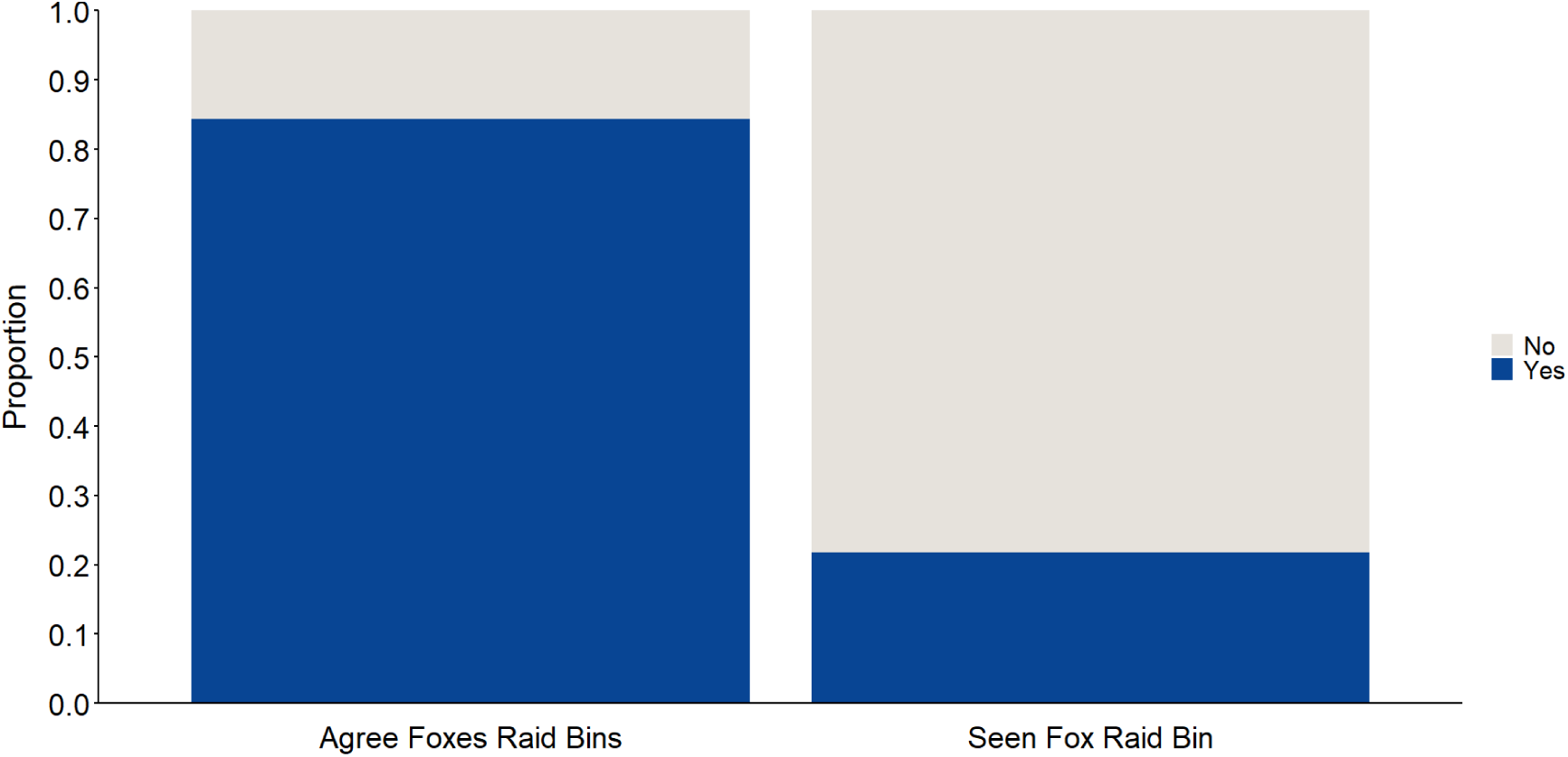
The proportion of people from the national public survey who agreed that foxes raid bins and have seen a fox raid a bin (N = 896 participants for agree foxes raid bins and 1,275 participants for seen fox raid bin).

Of the total 998 respondents from our national sample who reported that they had *not* witnessed a fox eating from a bin, 545 (54.6%) people agreed with the statement that foxes try to get food from peoples’ outdoor rubbish bins, while 114 (11.4%) and 339 (34%) people disagreed or were neutral, respectively. A Kruskal-Wallis test revealed that there was a significant difference in attitude scores towards foxes among participants in these three belief groups (Kruskal-Wallis: X^2^_2_= 24.52, p < 0.001). Post-hoc Dunn’s test revealed significant differences in two of the three pairwise comparisons. Specifically, respondents who agreed that foxes try to get food from people’s outdoor rubbish bins had significantly lower attitude scores (indicating less positive, or more negative attitudes towards foxes) than those who were neutral (Z = -3.47, p < 0.001) or those who disagreed (Z = -4.31, p < 0.001). There was no significant difference between respondents who disagreed with this statement and those who were neutral (Z = 1.88, p = 0.06).

In Hull specifically, 248 people responded to the public survey (Supplementary Materials), of which 42 (16.9%) people agreed with the statement that foxes raid bins, while 93 (37.5%) and 113 (45.6%) people disagreed or were neutral, respectively. In terms of respondents witnessing a local fox eating from a bin within the last year, 21 people (8.5%) said that they had, while the remaining 227 (91.5%) said that they had not. Similar to the national public questionnaire survey analysis, the proportion of respondents who agreed that foxes raid bins was significantly higher than the proportion who reported having personally witnessed fox bin-raiding in their local area within the last year (Two proportion Z test: X^2^_1_= 7.27, p = 0.007; **Figure 1.3**).

**Figure 1.3.**
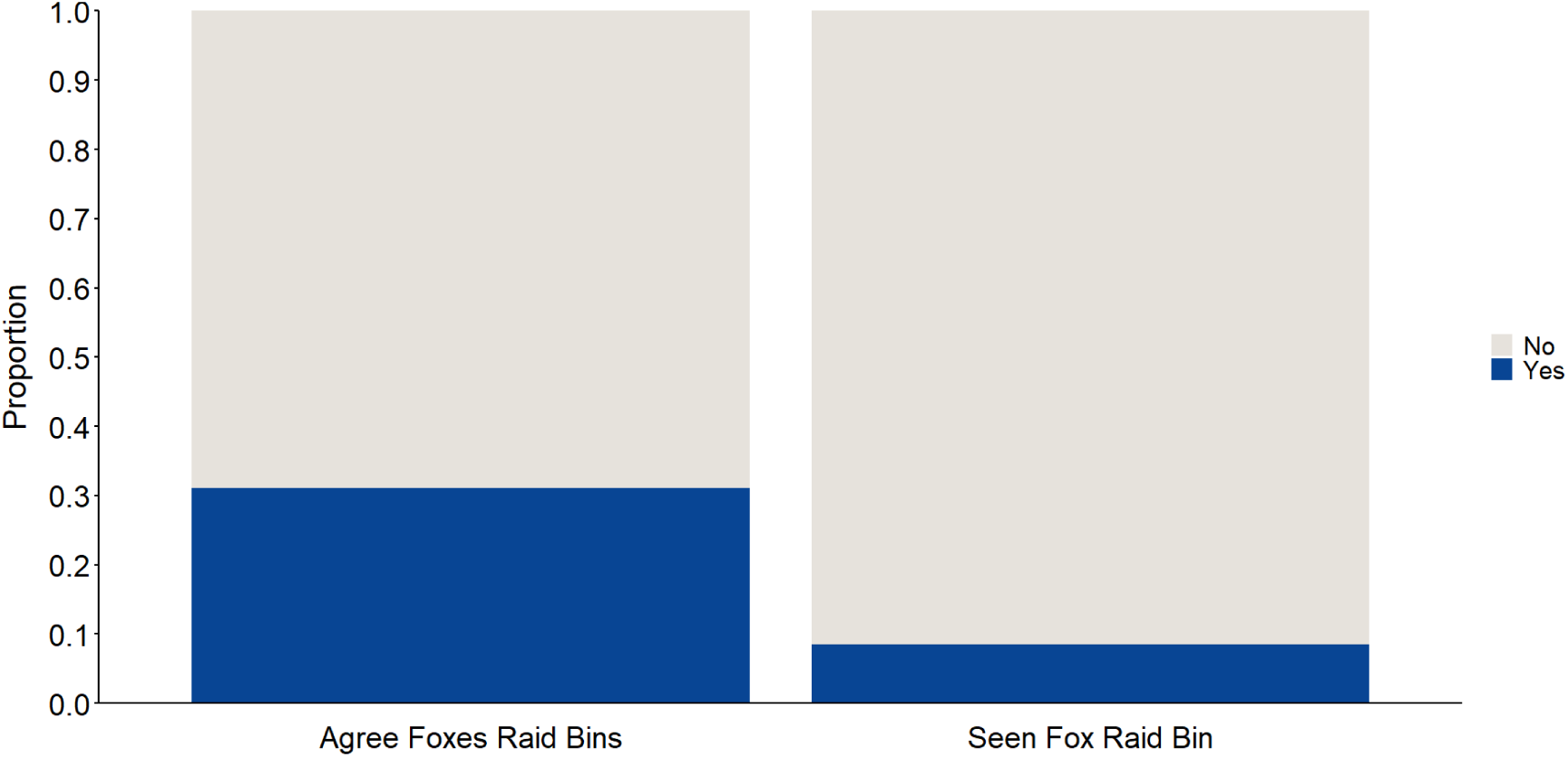
The proportion of people from the local public survey in Hull, UK, who agreed that foxes raid bins and have seen a fox raid a bin (N = 135 participants for agree foxes raid bins and 248 participants for seen fox raid bin).

Of the total 227 respondents from our local Hull sample who reported that they had *not* witnessed a local fox eating from a bin within the last year, 31 (13.7%) people agreed with the notion that foxes raid bins, 89 (39.2%) people disagreed, and 107 (47.1%) people were neutral. A Kruskal-Wallis test revealed that there was a significant difference in attitude scores towards foxes among participants in these three belief groups (Kruskal-Wallis: X^2^_2_= 25.14, p < 0.001). Post-hoc Dunn’s test revealed significant differences in two of the three pairwise comparisons. Specifically, respondents who agreed that foxes try to get food from people’s outdoor rubbish bins had significantly lower attitude scores (indicating less positive, or more negative attitudes towards foxes) than those who disagreed (Z = -4.34, p < 0.001). Additionally, respondents who disagreed with this statement had significantly higher attitude scores (more positive attitudes) than those who were neutral (Z = 3.96, p < 0.001). There was no significant difference between respondents who agreed with this statement and those who were neutral (Z = -1.66, p = 0.09).

### Ecological surveys on outdoor bin disturbances

A total of 4,239 bins were surveyed within the city of Hull, of which 2,866 (67.6%) were black/grey wheelie bins, 1,073 (25.3%) were brown wheelie bins, 133 (3.1%) were public street bins owned by the local council, 121 (2.9%) were dumpsters, 26 (0.6%) were green food caddies, 12 (0.3%) were wheelie bins of other colours, and 8 (0.2%) were bin bags placed on the ground (**Figure 1.4**).

**Figure 1.4.**
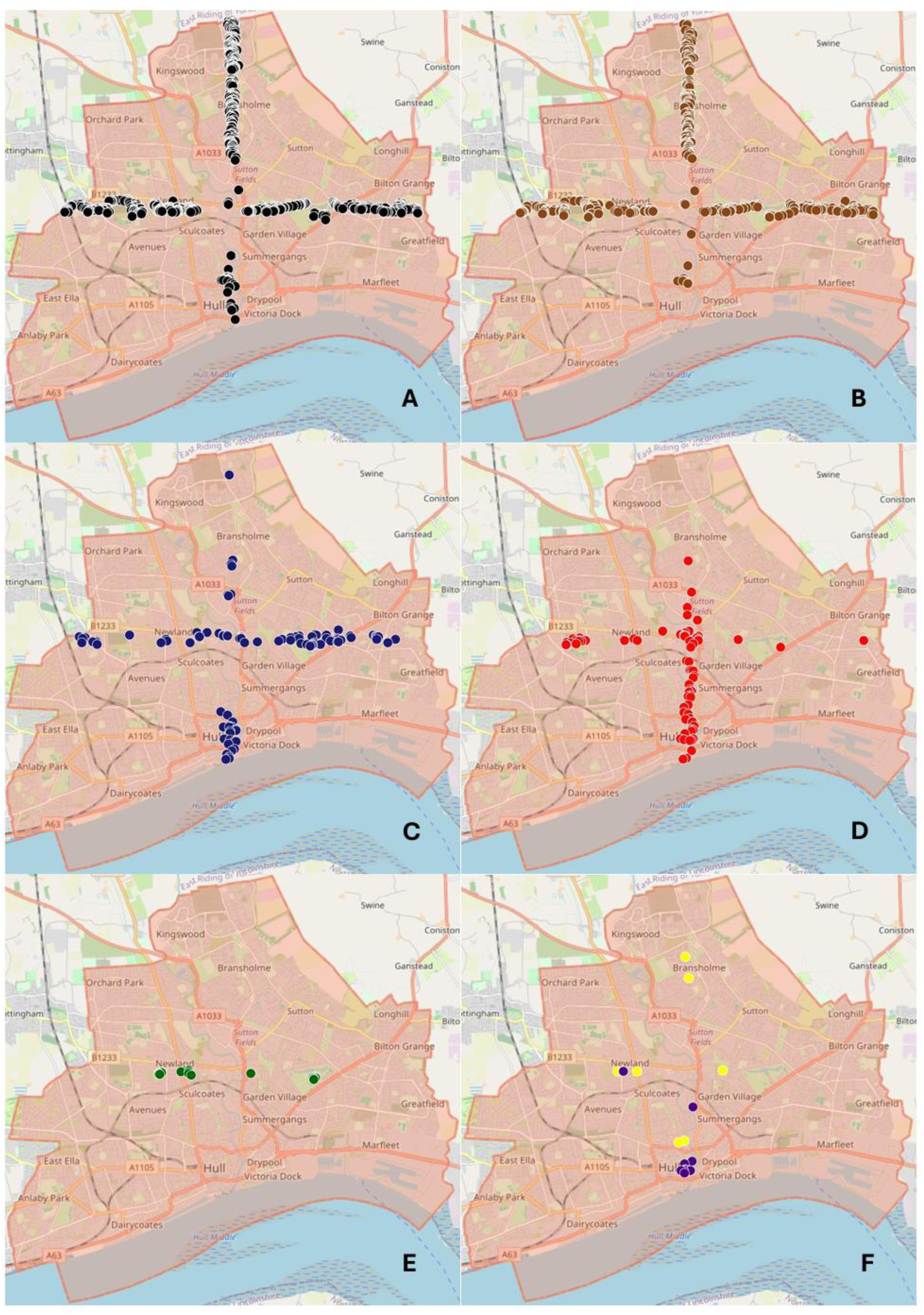
Ecological surveys to document outdoor bin disturbances across the city of Hull, UK. Including a) black wheelie bins (N = 2,866), b) brown wheelie bins (N = 1,073), c) public bins (N = 133), d) dumpsters (N = 121), e) food Caddies (N = 26), and f) other coloured wheelie bins in purple (N = 12) and bin bags in yellow (N = 8).

Using the conservative measure, only two (0.05%) bins were deemed likely to have been disturbed by an animal, including one grey wheelie bin and one bin bag. Using the less conservative measure, 19 (0.45%) bins were deemed to be potentially disturbed by an animal, including 12 grey wheelie bins, two brown wheelie bins, four bin bags and one public bin. In terms of accessibility, 488 (11.5%) bins were deemed to be easily accessible to foxes. All 19 bins identified in the less conservative measure were deemed to be easily accessible, though in two of these instances, accessibility was likely due to weather (i.e., high winds blowing bins over).

## Discussion

This study examined whether beliefs about wild red foxes reflect local observations of them, and whether public attitudes are associated with beliefs in the absence of local experience. Consistent with our hypotheses, people were more likely to believe that foxes raid bins compared to the number of people who had actually witnessed such behaviour locally. Moreover, among national and local Hull residents with no local experience observing bin-raiding foxes, beliefs about such behaviour were nevertheless associated with more negative attitudes towards the species. Objective bin surveys from Hull confirmed that instances of bin-raiding were rare in the area despite people from those neighbourhoods being more likely to believe foxes raided bins compared to the number of people who said they witnessed it firsthand. Together, these findings highlight the importance of taking into account other factors beyond local experiences when interpreting reports of human-wildlife conflict within a given area.

These findings are consistent with earlier ecological studies showing that bin-raiding is not a widespread behaviour in urban foxes and may, instead, reflect more localised instances from particular fox populations. Contesse et al. (2004), for example, found that only 1.3% of over 1,300 bin liners (i.e., the typical method of household waste disposal) in Zurich were damaged by foxes, or other animals, despite anthropogenic food making up a substantial part of fox diets. Moreover, our previous work found that instances of bin-raiding were primarily from the London area compared to other regions of the UK (Adaway et al., under review). While some UK studies have documented the presence of human-derived food in fox diets (Harris, 1981; Doncaster et al., 1990; Saunders et al., 1993; Williams et al., 2024; Fletcher et al., 2025), this does not necessarily mean such food is always taken from bins. Indeed, Doncaster et al. (1990) reported that 41% of their surveyed households regularly left food out for foxes and other wildlife, and other research has shown widespread wildlife feeding in urban settings (Baker et al., 2004; Baker & Harris, 2007; Gaston et al., 2007; Dorning & Harris, 2019; Scott et al., 2023). Thus, some anthropogenic items in fox diets may reflect deliberate provisioning rather than bin-raiding.

In the current study, 21.7% of respondents from the national public survey reported seeing a fox raiding bin, which is comparable to the 19.1% reported in a 1981 study conducted in Bristol, UK (Harris, 1981). Neither of these surveys specified a time frame for when these observations occurred, but only 8.5% of respondents in our local survey conducted in the city of Hull reported witnessing a fox raiding bin within the last year. While such differences may reflect regional or temporal variations in fox activity, they may also reflect the changes in human waste management practices. For instance, the Bristol study collected survey data between 1977 and 1978 (Harris, 1981), and in the 1980s, UK councils began replacing traditional dustbins with more secure wheelie bins (Chappells & Shove, 1999), which are harder for foxes to access. In later works, Harris and colleagues report that bin-raiding by foxes has declined since this change (Baker et al., 2004; Baker et al., 2006). Supporting this observation, all of the bins potentially raided by a fox in the current study’s bin surveys were deemed to be easily accessible, mainly due to improper waste management (e.g., bags left on the ground or bin lids wedged open). Remaining instances of bin-raiding incidents could, therefore, be shaped by human behaviour, such as poor waste storage. Future research should test this as well as other possibilities in other cities beyond Hull, particularly in London where reports of bin-raiding foxes are more prevalent.

Despite limited evidence for prolific bin-raiding behaviour by urban foxes, beliefs about conflict nevertheless persist among many people. Such persistent beliefs may be reinforced by other factors, such as portrayals of foxes in the media, particularly when articles depict foxes raiding bins or scavenging through waste (e.g., The Newsroom, 2013; Dubuis, 2014; Reynolds, 2025). Alternatively, perceptions may be socially transmitted, whereby people discuss experiences with foxes from other areas, regardless of direct local experience, thereby maintaining and circulating such beliefs (Dickman et al., 2014; Wilkinson et al., 2021; Songhurst, 2023). Further research is needed to test these and other possibilities.

In the UK, red foxes are not classified as vermin, but they have a long and complex history with people, both in rural and urban areas. Public beliefs about fox-related conflict, such as concerns about physical attacks in cities or predation game and wildlife in the countryside, frequently drive calls, and support for management or lethal control (Stewart & Cole, 2015; Bridge & Harris, 2020; Swan et al., 2020). While there are no laws specifically restricting the lethal management of foxes, broader legislation, including the Wildlife and Countryside Act (1981) and the Animal Welfare Act (2006), help to safeguard fox welfare. These laws, however, do not explicitly require that interventions be based on verified evidence of a physical conflict at local levels; in practice, generalised beliefs or reports of conflict in other areas can influence management decisions, sometimes leading to widespread persecution. Thus, the current study highlights a critical issue within this context: that local observations of wild animal behaviour do not always align with wider public beliefs. This mismatch illustrates a wider challenge for wildlife management across the UK and beyond. Indeed, management decisions that are guided primarily by perception rather than local empirical evidence risk being unnecessary, ethically problematic, and misaligned with the ecology of the species in question. Future research should therefore explore these discrepancies more broadly across taxa, including countries with large carnivores, urban-adapted species, and other wildlife commonly persecuted as threats or pests to society due to their perceived conflicts with people. Doing so will help to ensure ethical, evidence-based management is locally tailored to specific wildlife populations where instances of real (not perceived) conflict occur.

As discussed, the findings of this study could have important implications for the success of place-based conservation initiatives, such as urban rewilding, which can depend heavily on the public’s acceptance of wildlife (Pettorelli et al., 2022; Finnerty et al., 2025). Previous research shows that public intolerance of wildlife is associated with more negative attitudes towards them (Bruskotter et al., 2015; Puri et al., 2024). This is particularly important given that perceived local conflict and more negative attitudes have been linked to the use and support of lethal management of wildlife (Richard et al., 2005; Bruskotter et al., 2009; Hazzah et al., 2009; Echegaray & Vilà, 2010; Engel et al., 2017; Ballejo et al., 2020; Kimmig et al., 2020). The current study found that respondents who believed that bin-raiding occurred, despite not experiencing it locally themselves, held significantly more negative attitudes towards foxes compared to those who did not hold the belief. It is unclear whether negative attitudes predispose individuals to believe conflict occurs (e.g., confirmation bias; Wieczorek Hudenko, 2012) or whether beliefs about conflict influence attitudes. Nevertheless, the observed association highlights a key barrier to coexistence: Even if negative attitudes are formed first, beliefs about wildlife causing conflict could still justify or intensify support for lethal control measures, thereby influencing wildlife management outcomes. If decisions are guided by broader public attitudes and beliefs rather than direct local objective evidence, they risk being disproportionate to the actual conflict, causing unnecessary harm to wildlife populations. Incorporating strategies such as effective environmental messaging to target non-local factors may, therefore, be essential for overcoming such biases to promote tolerance and more ethical, evidence-based approaches to conservation management.

## Conclusion

This study demonstrates a clear discrepancy between public beliefs about bin-raiding in wild red foxes and the actual frequency of such behaviour observed by people within their local area. Agreement with the notion of bin-raiding foxes among respondents who had not witnessed it locally was associated with more negative attitudes towards the species. Such findings highlight that public beliefs about foxes’ bin-raiding can be maintained even in the absence of local experience, consistent with the notion that factors beyond direct observation influence attitudes towards wildlife, particularly socio-cultural narratives. Such mismatches have important implications for wildlife management and conservation, as decisions driven primarily by subjective reports of conflict are at risk of being disproportionate, ethically questionable, and misaligned with more nuanced, localised animal behaviour. Promoting human-wildlife coexistence may require strategies that address not only local experiences, but also the broader social and informational factors influencing public attitudes and beliefs to ensure that management actions are both evidence-based and locally appropriate. Doing so may be crucial to the success of place-based conservation initiatives, such as urban rewilding, where local tolerance and coexistence with species is necessary.

## Notes

### Competing Interest Statement

The authors have declared no competing interest.

