## Supplementary materials for "Beliefs about a widely popularised human-wildlife conflict are largely unreflective of local experiences and ecological data in wild red foxes, *Vulpes vulpes*"

**Local public survey questions sent to residents of the HU88 and HU54 postal areas of the City of Hull:**

**Part A** Question 1:

Please complete the following two-part survey. Your participation is entirely voluntary, and your responses will remain anonymous. It takes approximately 5 minutes to complete, but there is no time limit so please take however long you need. We greatly appreciate your time and thoughts!

Please note that participation in the study requires that you consent to the following statements. If you decide to take part, please click “I wish to proceedˮ. If you decide not to take part, please exit the survey before submitting your responses.

- I confirm that I have read the “Participant Information Sheetˮ that explains the study.
- I have had the opportunity to consider the information and have been given contact details of the researcher to ask questions and discuss the study.
- Any questions or concerns I have about the study have been answered satisfactorily.
- I understand that my participation is voluntary and that I am free to withdraw at any time during the survey without giving any reason.
- I understand that once I have completed the survey, I cannot withdraw my anonymised data.
- I understand that the research data, which are not linked to me, will be retained by the researchers, and may be shared with others and publicly disseminated to support other research in the future.
- I agree to take part in the study.
- I wish to proceed
- I do not wish to proceed

Question 2:

What is your current age in years?

- 18-24
- 25-34
- 35-44
- 45-54
- 55-64
- 65+
- Prefer not to say

Question 3:

What is your gender?

- Male
- Female
- Nonbinary
- Other
- Prefer not to say

Question 4:

Do you currently live in the city of Hull?

- Yes
- No

Question 5:

If you live in the city of Hull, what is the first half of your postcode (e.g., HU8, HU12)?

*Open response*

Question 6:

Does your current place of residence have an outdoor garden?

- Yes
- No

Question 7:

At your current place of residence, how often do you put food in your outdoor bin (e.g., wheelie bin or food waste bin)?

- I do not have an outdoor bin
- Never
- Less than once a week
- 1 to 3 times per week
- More than 3 times per week

Question 8:

From the image below, please select the letter shown below the image of a fox

- A
- B
- C

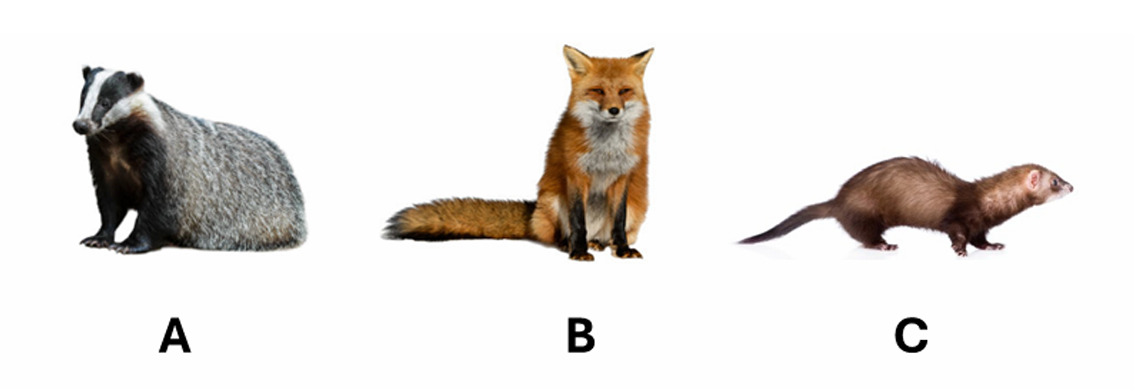

Question 9:

Within the last year have you seen (in person or on a camera) a fox around your current place of residence (e.g., in your garden or on the residential street outside your home)?

- Yes
- No

Question 10:

At your current place of residence, how often do you intentionally leave food outside for foxes?

- Never
- Less than once a week
- 1 to 3 times per week
- More than 3 times per week

Question 11:

At your current place of residence, how often do you intentionally leave food outside for other wildlife (e.g., birds or hedgehogs)?

- Never
- Less than once a week
- 1 to 3 times per week
- More than 3 times per week

Question 12:

At your current place of residence, have you seen (in person or on camera) a fox eating rubbish from your own OR a neighbour's outdoor bin within the last year?

- Yes
- No
- Neither myself or my neighbours have outdoor bins

Question 13:

From the selection below, please select the type(s) of bin belonging to yourself OR your neighbour that you have seen a fox raid within the last year (i.e., colour of bins not important, e.g., please select black wheelie bin for brown wheelie bins). Check all that apply.

- A
- B
- C
- D
- E
- Other design of bin not pictured

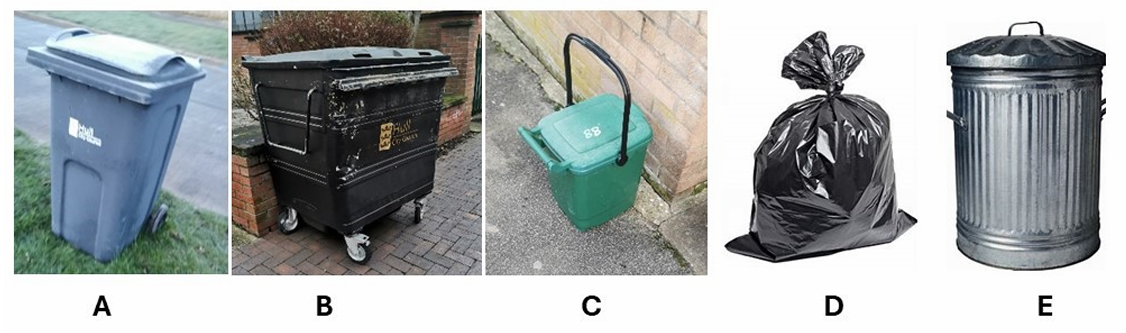

Question 13:

If you selected other, please describe the other type(s) of bin to the best of your ability (e.g., rough height, material and structure).

*Open response*

Question 14:

Select from the list below the season(s) in which you saw fox eating rubbish from your own OR a neighbour's outdoor bin during the last year. Check all that apply.

- Spring
- Summer
- Autumn
- Winter
- I do not know

Question 15:

Was the lid to your own OR your neighbour's bin(s) left open at the time when you saw a fox raiding it within the last year (i.e., lid wedged open with contents)?

- Yes
- No
- On some occasions but not every time
- I do not know

Question 16:

At your current place of residence, have you ever tried to harm or kill a fox (e.g., poison or hunt them)?

- Yes
- No

Question 17:

At your current place of residence, have you ever contacted a pest control service to resolve an issue you had with foxes?

- Yes
- No

Question 18:

Within the last year have you seen a fox within your neighbourhood?

- Yes
- No

Question 19:

Within your neighbourhood, have you ever seen (in person or on camera) a fox eating rubbish from a public bin owned by your local council within the last year?

- Yes
- No
- The local council does not have bins in my area

Question 20:

From the selection below, please select the type(s) of public bin that you have seen a fox raid within the last year (i.e., colour of bins not important). Check all that apply.

- A
- B
- C
- D
- E
- Other design of bin not pictured

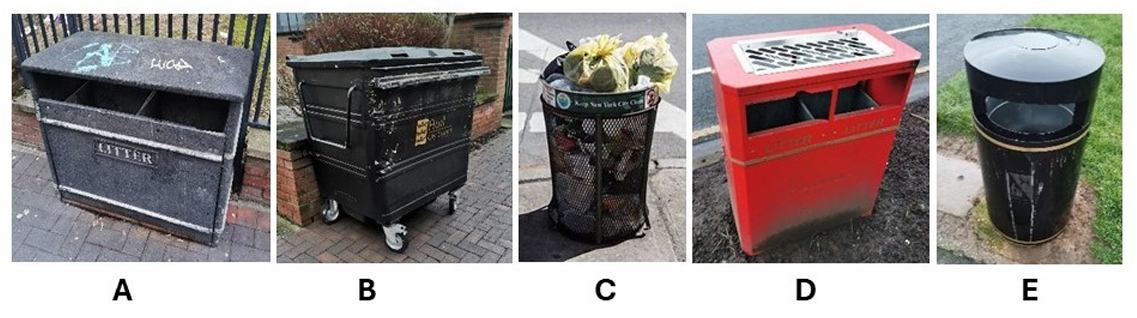

Question 21:

If you selected other, please describe the other type of public bin(s) to the best of your ability (e.g., rough height, material and structure).

*Open response*

Question 22:

Select from the list below the season(s) in which you saw fox eating rubbish from a public bin owned by your local council during the last year. Check all that apply.

- Spring
- Summer
- Autumn
- Winter
- I do not know

Question 23:

Was the public bin(s) overflowing at the time when you saw a fox raiding it within the last year (i.e., rubbished filled to the top)?

- Yes
- No
- On some occasions but not every time
- I do not know

**Part B**

Please click next to continue to part B of the survey.

For the following questions you will be asked to what extent you agree with 20 statements regarding foxes.

Question 1:

The number of foxes should be controlled by human shooting or other forms of control (e.g., poisoning).

- 1 (Strongly disagree)
- 2
- 3
- 4 (Neither agree nor disagree)
- 5
- 6
- 7 (Strongly agree)

Question 2:

It is acceptable for people to harm or kill wild foxes.

- 1 (Strongly disagree)
- 2
- 3
- 4 (Neither agree nor disagree)
- 5
- 6
- 7 (Strongly agree)

Question 3:

Foxes are potential carriers of diseases and shouldnʼt be around people.

- 1 (Strongly disagree)
- 2
- 3
- 4 (Neither agree nor disagree)
- 5
- 6
- 7 (Strongly agree)

Question 4:

Because of the presence of wild foxes, I would be scared to walk alone outdoors.

- 1 (Strongly disagree)
- 2
- 3
- 4 (Neither agree nor disagree)
- 5
- 6
- 7 (Strongly agree)

Question 5:

The presence of wild foxes would negatively affect my leisure activities.

- 1 (Strongly disagree)
- 2
- 3
- 4 (Neither agree nor disagree)
- 5
- 6
- 7 (Strongly agree)

Question 6:

Foxes try to retrieve litter and other discarded food containers shortly after discovering them.

- 1 (Strongly disagree)
- 2
- 3
- 4 (Neither agree nor disagree)
- 5
- 6
- 7 (Strongly agree)

Question 7:

Foxes try to get food from peoplesʼ outdoor rubbish bins.

- 1 (Strongly disagree)
- 2
- 3
- 4 (Neither agree nor disagree)
- 5
- 6
- 7 (Strongly agree)

Question 8:

Foxes are a “nuisanceˮ in my everyday life.

- 1 (Strongly disagree)
- 2
- 3
- 4 (Neither agree nor disagree)
- 5
- 6
- 7 (Strongly agree)

Question 9:

Foxes are dangerous for children.

- 1 (Strongly disagree)
- 2
- 3
- 4 (Neither agree nor disagree)
- 5
- 6
- 7 (Strongly agree)

Question 10:

I consider foxes in urban environments a pest.

- 1 (Strongly disagree)
- 2
- 3
- 4 (Neither agree nor disagree)
- 5
- 6
- 7 (Strongly agree)

Question 11:

Wild foxes should only live in nature reserves and other protected areas.

- 1 (Strongly disagree)
- 2
- 3
- 4 (Neither agree nor disagree)
- 5
- 6
- 7 (Strongly agree)

Question 12:

I enjoy seeing foxes.

- 1 (Strongly disagree)
- 2
- 3
- 4 (Neither agree nor disagree)
- 5
- 6
- 7 (Strongly agree)

Question 13:

I would be pleased having a fox in my garden/living environment.

- 1 (Strongly disagree)
- 2
- 3
- 4 (Neither agree nor disagree)
- 5
- 6
- 7 (Strongly agree)

Question 14:

It is acceptable to see a fox in my own garden or neighbourhood.

- 1 (Strongly disagree)
- 2
- 3
- 4 (Neither agree nor disagree)
- 5
- 6
- 7 (Strongly agree)

Question 15:

It is acceptable for more foxes to live in my neighbourhood (i.e., fox population growth).

- 1 (Strongly disagree)
- 2
- 3
- 4 (Neither agree nor disagree)
- 5
- 6
- 7 (Strongly agree)

Question 16:

Foxes are part of nature. They belong to our environment and should be accepted around humans.

- 1 (Strongly disagree)
- 2
- 3
- 4 (Neither agree nor disagree)
- 5
- 6
- 7 (Strongly agree)

Question 17:

Wild foxes have, like other animals, a right to live in the UK.

- 1 (Strongly disagree)
- 2
- 3
- 4 (Neither agree nor disagree)
- 5
- 6
- 7 (Strongly agree)

Question 18:

The presence of wild foxes increases the value of a landscape, whether I get to see them or not.

- 1 (Strongly disagree)
- 2
- 3
- 4 (Neither agree nor disagree)
- 5
- 6
- 7 (Strongly agree)

Question 19:

For me, it is important to protect wild fox populations also for future generations.

- 1 (Strongly disagree)
- 2
- 3
- 4 (Neither agree nor disagree)
- 5
- 6
- 7 (Strongly agree)

Question 20:

Only foxes that cause problems and damages should be controlled through scaring, capturing, relocating or shooting.

- 1 (Strongly disagree)
- 2
- 3
- 4 (Neither agree nor disagree)
- 5
- 6
- 7 (Strongly agree)

Question 21:

Since 2021, scientists from the University of Hull have been running a citizen science programme called The British Carnivore Project, where they have been studying the impact of urbanization and climate change on the cognition and behaviour of wild free-ranging foxes and badgers. Please click all items that best describe your involvement in that programme.

- I have read about the project online.
- I was copied into emails about the project
- I gave permission to access the land
- I helped collect data or monitor a trail camera
- I read through the information about the projectʼs goals and current research findings
- I had no involvement in the study

END OF SURVEY - Please click submit to save your responses.

Thank you for completing the survey.

**Table S4.1**. Cohen’s kappa inter-observer reliability test for whether bins were easily accessible to animals by K.A and independent coder (C. B) who coded 100% of the videos.

|  | Count |
| --- | --- |
| Accessible (agree) | 427 |
| Not accessible (agree) | 3737 |
| Disagreements | 75 |
| Total | 4239 |
| IOR result | K = 0.909 |

**Table S4.2**. Cohen’s kappa inter-observer reliability test for whether bins were “highly likely to have been disturbed by an animal” (i.e., the conservative measure) that was potentially a fox, by K.A and an independent coder (K. S.) who coded 100% of the videos.

|  | Count |
| --- | --- |
| Highly likely disturbed by an animal (agree) | 2 |
| Not highly likely to have been disturbed by an animal (agree) | 4237 |
| Disagreements | 0 |
| Total | 4239 |
| IOR result | K = 1 |

**Table S4.3**. Cohen’s kappa inter-observer reliability test for whether bins were “potentially disturbed by an animal” (i.e., the non-conservative measure), by K.A and an independent coder (D. J) who coded 50 of the videos, 1 = potentially disturbed, 0 = not disturbed (k = 0.63).

| Bin Number | K. A. | D. J. (Independent coder). |
| --- | --- | --- |
| 1 | 0 | 0 |
| 2 | 0 | 0 |
| 3 | 0 | 0 |
| 4 | 0 | 0 |
| 5 | 1 | 1 |
| 6 | 1 | 0 |
| 7 | 0 | 0 |
| 8 | 1 | 0 |
| 9 | 0 | 0 |
| 10 | 0 | 0 |
| 11 | 0 | 0 |
| 12 | 0 | 0 |
| 13 | 0 | 0 |
| 14 | 1 | 1 |
| 15 | 0 | 0 |
| 16 | 1 | 0 |
| 17 | 0 | 0 |
| 18 | 0 | 0 |
| 19 | 0 | 0 |
| 20 | 0 | 0 |
| 21 | 1 | 0 |
| 22 | 0 | 0 |
| 23 | 1 | 1 |
| 24 | 1 | 1 |
| 25 | 0 | 0 |
| 26 | 1 | 1 |
| 27 | 1 | 0 |
| 28 | 1 | 1 |
| 29 | 1 | 0 |
| 30 | 0 | 0 |
| 31 | 1 | 1 |
| 32 | 0 | 0 |
| 33 | 0 | 0 |
| 34 | 1 | 1 |
| 35 | 0 | 0 |
| 36 | 0 | 0 |
| 37 | 1 | 0 |
| 38 | 1 | 1 |
| 39 | 0 | 0 |
| 40 | 1 | 1 |
| 41 | 1 | 1 |
| 42 | 0 | 0 |
| 43 | 0 | 0 |
| 44 | 0 | 0 |
| 45 | 0 | 0 |
| 46 | 0 | 0 |
| 47 | 0 | 0 |
| 48 | 1 | 0 |
| 49 | 0 | 0 |
| 50 | 0 | 0 |

**Note**: In this instance, both coders agreed on eleven instances where a bin was “potentially disturbed by an animal” (i.e., the non-conservative measure), and 31 instances where the bins were not potentially disturbed by an animal. The remaining eight differences arose due to K. A. identifying eight bins as potentially disturbed that D. J. did not. The differences between K. A’s and D. J’s coding is therefore due to the coding of K. A. been extremely cautious as to not exclude any instances where it is possible an animal disturbed the scene and are therefore still included within the study.

**Supplementary discussion**

**Note on local public survey’s low response rates**

Overall, the local survey had a low response rate of 3.07%. The survey was advertised through unaddressed leaflets delivered to residential homes using a postal service, and no incentives were offered for participation. Low response rates are expected when using flyer-based recruitment methods (Al-Muhanna et al., 2023). However, such rates can still introduce the possibility of non-response bias (i.e., where those who respond may differ systematically from those who do not). Nevertheless, it is worth noting that the results closely aligned with those of the national survey, which was distributed via a commercial survey company and offered participants a £5 voucher incentive. This consistency suggests that, despite potential sampling biases, the local survey may still reflect meaningful and representative patterns in public attitudes.
